# Virological and antigenic characteristics of SARS-CoV-2 variants LF.7.2.1, NP.1, and LP.8.1

**DOI:** 10.1101/2024.12.27.630350

**Authors:** Jingyi Liu, Yuanling Yu, Sijie Yang, Fanchong Jian, Weiliang Song, Lingling Yu, Fei Shao, Yunlong Cao

## Abstract

XEC and KP.3.1.1 have surpassed KP.3 to become the globally dominant lineages due to their unique NTD mutations. However, several emerging JN.1 sublineages, such as LF.7.2.1, MC.10.1, NP.1, and, especially, LP.8.1, have demonstrated superior growth advantages compared to XEC. It is critical to access the virological and antigenic characteristics of these emerging SARS-CoV-2 variants. Here, we found that LF.7.2.1 is significantly more immune invasive than XEC, primarily due to the A475V mutation, which enabled the evasion of Class 1 neutralizing antibodies. However, LF.7.2.1’s weak ACE2 binding affinity substantially impaired its fitness. Likewise, MC.10.1 and NP.1 exhibited strong antibody immune evasion due to the A435S mutation, but their limited ACE2 engagement efficiency restricted their growth advantage, suggesting that A435S may regulate the Spike conformation, similar to the NTD glycosylation mutations found in KP.3.1.1 and XEC. Most importantly, we found that LP.8.1 showed comparable humoral immune evasion to XEC but demonstrated much increased ACE2 engagement efficiency, supporting its rapid growth. These findings highlight the trade-off between immune evasion and ACE2 engagement efficiency in SARS-CoV-2 evolution, and underscore the importance of monitoring LP.8.1 and its descend lineages.

## Main

Recently, XEC and KP.3.1.1 have surpassed KP.3 to become the globally dominant lineages due to the unique N-terminal domain (NTD) mutations, including S31del in KP.3.1.1 and T22N and F59S in XEC^1-7^. However, several sublineages of JN.1 are increasingly out-competing XEC and KP.3.1.1, exhibiting superior growth advantages. For example, LF.7.2.1, MC.10.1, NP.1, and most importantly, LP.8.1 (Figures 1A and 1B). Notably, LF.7.2.1 contains an additional A475V mutation compared to LF.7, which carries the S31P, K182R, R190S, and K444R mutations on Spike, and has rapidly spread from Qatar to the Middle East and Europe. MC.10.1, a KP.3.1.1 subvariant with a rare spike mutation A435S, shows a slightly higher growth advantage than KP.3.1.1^*^. Meanwhile, NP.1, with an additional S446N on the receptor-binding domain (RBD), exhibits enhanced growth advantage and has spread rapidly in Canada. LP.8, a subvariant of KP.1.1, carries S31del, F186L, Q493E, and H445R on Spike. Importantly, LP.8.1, with an additional R190S, has surged rapidly in America, exhibiting the highest growth advantage among circulating variants (Figure 1B). These novel RBD and NTD mutations have given rise to fast-evolving variants, underscoring the urgent need to evaluate their virological and antigenic characteristics for future preparedness.

**Figure 1.**
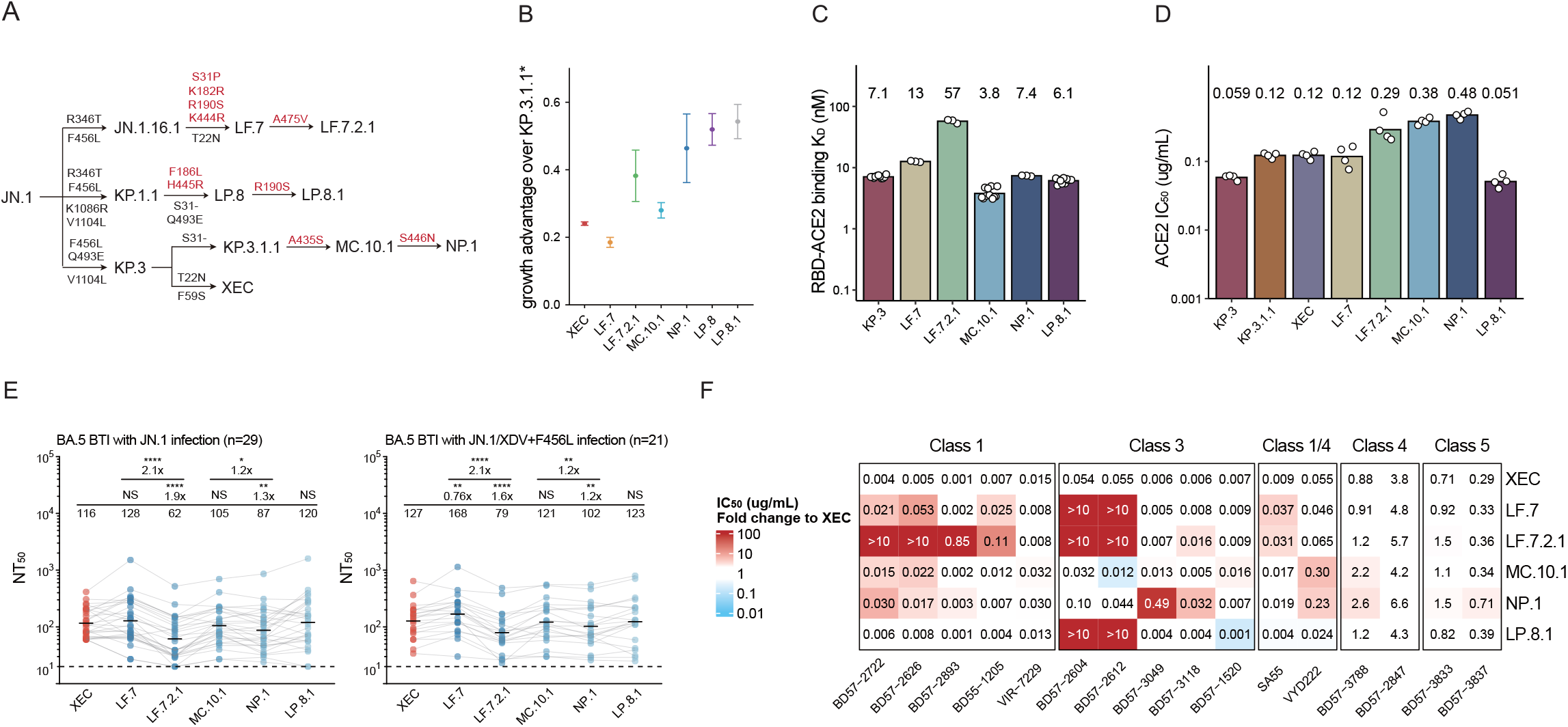
ACE2 engagement and antibody evasion characteristics of LF.7.2.1, MC.10.1, NP.1, and LP.8.1. (A) This schematic shows the evolution of the Spike glycoprotein in prevalent SARS-CoV-2 variants. The mutations highlighted in red are specific to these variants, in addition to those shared with KP.3.1.1 and XEC. (B) This chart shows the relative growth advantage of XEC, LF.7, LF.7.2.1, MC.10.1, NP.1, LP.8, LP.8.1, compared to KP.3.1.1^*^. The relative growth advantage was calculated from daily sequence data sourced from the GISAID database. ^*^Indicates all sublineages. (C) This chart shows the binding affinity of KP.3, LF.7, LF.7.2.1, MC.10.1, NP.1, and LP.8.1 RBD proteins to human ACE2, established by SPR. Each circle indicates a replicate. Geometric mean K_D_ values (nM) are displayed above each bar. K_D_=dissociation constant. (D) This barplot shows the IC_50_ values of soluble human ACE2 against KP.3, KP.3.1.1, XEC, LF.7, LF.7.2.1, MC.10.1, NP.1, and LP.8.1 pseudoviruses. Each circle indicates a replicate. IC_50_ values (μg/mL) are displayed above each bar. IC_50_=50% inhibitory concentration. (E) These graphs show the NT_50_ of convalescent plasma from individuals reinfected with JN.1 after BA.5 or BF.7 breakthrough infection (n=29) and those reinfected with JN.1 or XDV with F456L after BA.5 or BF.7 breakthrough infection (n=21). Plasma source cohorts and corresponding number of samples are labeled above each panel. Dashed line indicates limit of detection (NT_50_=10). Geometric mean titres are labelled above each group, with fold changes and statistical significance indicated above the geometric mean titre labels. Wilcoxon rank-sum tests are used to determine the p-values. ^*^p<0.1, ^**^p<0.01, ^****^p<0.0001, NS=non-significant (p>0.05). NT_50_=50% neutralising titre. (F) The table shows IC_50_ values for a panel of monoclonal neutralising antibodies targeting RBD epitopes against XEC, LF.7, LF.7.2.1, MC.10.1, NP.1, and LP.8.1 variants. The values within the table are IC_50_ values (µg/mL), while the background color indicates the fold-change in IC_50_ relative to XEC. The color gradient bar to the left represents the magnitude of IC_50_ fold-change, with red indicating IC_50_ significantly higher than XEC and blue indicating lower. Antibodies tested are listed below the table, grouped by their target classes (Class 1, Class 3, Class 1/4, Class 4, and Class 5), shown above the table. The tested variants are indicated on the right.

We first constructed the recombinant RBD subunit of KP.3, LF.7, LF.7.2.1, MC.10.1, NP.1, and LP.8.1 and assessed their binding affinity to human ACE2 (hACE2) using surface plasmon resonance (SPR) (Figure 1C). The results show that the A475V mutation in LF.7.2.1 greatly reduced RBD-ACE2 binding affinity compared to LF.7. In contrast, the binding affinities of LF.7, LP.8.1, and NP.1, carrying K444R, H445R, and S446N, respectively, did not exhibit substantial changes. The A435S mutation in MC.10.1 resulted in a slight increase in binding affinity compared to KP.3.

Given that RBD-ACE2 binding affinity alone does not fully reflect the effect of mutations on other regions or the Spike trimer conformation, we constructed Spike-pseudotyped vesicular stomatitis virus (VSV) and assayed the efficiency of soluble hACE2 to inhibit viral entry, which indicates the ACE2 binding strength and receptor engagement efficiency of the variants Spike proteins (Figure 1D). Consistent with previous findings, mutations on the NTD of KP.3.1.1 and XEC decreased the engagement efficiency compared to KP.3^1-4^. In concordance with the SPR results, the A475V mutation in LF.7.2.1 greatly reduced the engagement efficiency compared to LF.7. Surprisingly, we found that the A435S mutation reduced ACE2 engagement efficiency in MC.10.1 and NP.1 compared to KP.3.1.1, possibly by altering the conformation of the RBD. Most importantly, LP.8.1 enhances ACE2 engagement efficiency to a level similar to KP.3, potentially contributing to its highest growth advantage.

Next, we evaluated the humoral immune evasion by using convalescent plasma and a panel of RBD-targeting neutralizing monoclonal antibodies (mAbs) in pseudovirus neutralization assays (Figures 1E and 1F). The plasma used in this study was obtained from two cohorts of individuals who received 2–3 doses of inactivated SARS-CoV-2 vaccines and subsequently experienced BA.5 or BF.7 breakthrough infections, with one cohort reinfected by JN.1 (n=29) and the other by JN.1 or XDV with F456L (n=21), as previously described^1^. Compared to KP.3, LF.7 showed similar or slightly reduced plasma immune evasion, while LF.7.2.1 significantly increased it, making it currently the most immune-evasive variant. This enhancement is attributed to the A475V mutation, which significantly strengthened resistance to Class 1 antibodies, consistent with previous studies (Figure 1F)^8-10^. MC.10.1 exhibited high immune evasion similar to XEC, with the RBD mutation A435S reducing the neutralization potency of Class 1 and Class 1/4 antibodies. Since site 435 is not located on the epitope of Class 1 antibodies, this suggests that A435S may enhance immune evasion by modulating RBD conformation, a mechanism similar to that of the NTD mutations in KP.3.1.1 and XEC. Additionally, NP.1, which carries an additional S446N, further enhanced immune evasion by improving its ability to escape Class 3 antibodies. Importantly, LP.8.1 maintained a high level of humoral immune evasion similar to XEC, while additionally escaping recognition by certain Class 3 antibodies.

In summary, LF.7.2.1 is the most immune evasive variant, but its relatively weak ACE2 binding affinity and engagement efficiency explains why it does not exhibit the highest growth advantage despite its strong immune evasion. Similarly, MC.10.1 and NP.1 show strong immune evasion, but their limited ACE2 engagement efficiency restricts the growth advantage. Importantly, we find that LP.8.1 exhibits exceptionally high ACE2 engagement efficiency as while as high immune evasion similar to XEC. The emergence of KP.3.1.1 and XEC involved a trade-off, with enhanced immune evasion achieved at the expense of ACE2 engagement efficiency, thereby affecting their fitness. In contrast, LP.8.1 has found a way to preserve ACE2 engagement efficiency similar to KP.3, while achieving immune evasion capabilities akin to XEC. LP.8.1’s current dominant growth advantage underscores the urgent need for contemporary variants to regain ACE2 engagement efficiency. These findings highlight the strong trade-off between immune evasion and ACE2 engagement efficiency in SARS-CoV-2 evolution, underscoring the importance of monitoring LP.8.1, especially after its recent convergent acquisition of the A475V mutation.

## Acknowledgments

We extend our gratitude to the scientific community for their continued efforts in monitoring SARS-CoV-2 variants, as well as to all volunteers who contributed blood samples for this study. This work was supported by National Science and Technology Major Project (2022ZD0115002), and the National Natural Science Foundation of China (32222030, 2023011477).

## Author Contributions

Y.C. designed and supervised the study. J.L. and Y.C. wrote the manuscript with inputs from all authors. J.L., S.Y., F.J., and W.S. performed sequence analysis, illustration, and figure preparation.

Y.Y. constructed pseudoviruses. L.Y., and F.S. processed the plasma samples and performed the pseudovirus neutralisation assays. F.J. and Y.C. analyzed the neutralisation data.

## Methods

### Growth Advantage Calculation

The algorithm for calculating growth advantage was adapted from Chen et al^11^. The relative growth advantage of each variant was calculated using a logistic regression model, which was fit to the daily frequency of samples from the GISAID database^12^. The model estimated the logistic growth rate (a) and midpoint (t0) of the sigmoid curve for each strain. The growth advantage was determined as e^a × g^ −1, where g was set as 7 days, the generation time. Confidence intervals were computed with α = 0.95. The total number of sequences for the target variant and the reference strains were used as the background to estimate the relative growth advantage.

### Surface Plasmon Resonance

SPR experiments were performed using Biacore 8K Evaluation Software 3.0 (Cytiva). After a 1-hour incubation at 37°C with 5% CO2, digested human ACE2 was immobilized on Sensor Chip CM5 (Cytiva). Purified RBD samples (His Tag, Sino Biological) of various concentrations (6.25, 12.5, 25, 50, and 100 nM) were serially diluted and injected onto the sensor chips. Data acquisition was carried out at room temperature using the Biacore 8K, and the binding kinetics were analyzed with a 1:1 binding model in the software. Each variant was tested in three independent replicates to ensure the reliability of the results.

### Patient recruitment and plasma isolation

Blood samples were collected from volunteers who had been reinfected with the SARS-CoV-2 Omicron BTI variant, following a research protocol approved by the Ethics Committees of Beijing Ditan Hospital Capital Medical University (Ethics Committee Archiving No. LL-2021-024-02), the Tianjin Municipal Health Commission, and the ethics committee of Tianjin First Central Hospital (Ethics Committee Archiving No. 2022N045KY). Informed consent was obtained from all participants in accordance with the Declaration of Helsinki. Each participant provided written consent for the collection, storage, and use of their clinical samples for research purposes, as well as for the publication of the data generated from this study.

The majority of patients were initially infected in December 2022 in Beijing and Tianjin, primarily involving the BA.5^*^ variants. Between December 1, 2022, and February 1, 2023, over 98% of sequenced samples were identified as BA.5/BF.7 variants, primarily including subtypes BA.5.2.48, BF.7.14, etc. Patients in the JN.1 reinfection cohort had their first breakthrough infection after receiving two to three doses of inactivated vaccine, followed by reinfection between February and March 2024. And the volunteers in the JN.1/XDV + F456L reinfection cohort, who experienced an initial outbreak of infection after receiving three doses of inactivated vaccine, were reinfected in July-August 2024, during which more than 97% of the sequenced samples were identified as JN.1 + F456L or XDV + F456L. The infections are confirmed by polymerase chain reaction (PCR) or antigen assay.

Whole blood samples were diluted 1:1 in PBS + 2% FBS and then subjected to gradient centrifugation using Ficoll (Cytiva, 17-1440-03) to isolate plasma and PBMCs. After centrifugation, the plasma was collected from the upper layer. The resulting plasma samples were collected, aliquoted, and stored at -20°C or lower. Heat inactivation was performed before further experimental use.

### Pseudovirus neutralisation assay

Vesicular stomatitis virus (VSV) pseudovirus packaging system was used to package SARS-CoV-2 variant spike pseudovirus. The G^*^ΔG-VSV virus (VSV G pseudotyped virus, Kerafast) was used to infect 293T cells (American Type Culture Collection [ATCC], CRL-3216), and the Spike-expressing plasmid was used for transfection at the same time. After cell culture, the pseudovirus in the supernatant was collected, filtered, aliquoted, and frozen at −80 °C for further use.

Monoclonal antibodies, human ACE2-Fc, and plasma samples were serially diluted in culture media, mixed with the pseudovirus in 96-well plates, and incubated at 5% CO_2_ and 37°C for 1 hour. Plasma samples were assayed at a 1:10 starting dilution and three additional tenfold serial dilutions. Anti-S-RBD antibody and ACE2-Fc were both tested at a 0.1mg/mL starting concentration and in additional fivefold serial dilutions. Then, digested Huh-7 cells (Japanese Collection of Research Bioresources [JCRB], 0403) or Vero cells (American Type Culture Collection [ATCC], CCL-81) were seeded. We used Huh-7 cells in the plasma neutralisation experiments to maintain consistency with previous experimental data. Following 24 hours incubation, the supernatant was removed, and D-luciferin reagent (PerkinElmer, 6066769) was added to initiate the reaction in the dark for 2 minutes. Luminescence was then measured using a microplate spectrophotometer (PerkinElmer, HH3400). The IC50 value was calculated using a four-parameter logistic regression model.

## Supplementary Tables

**Table S1.**
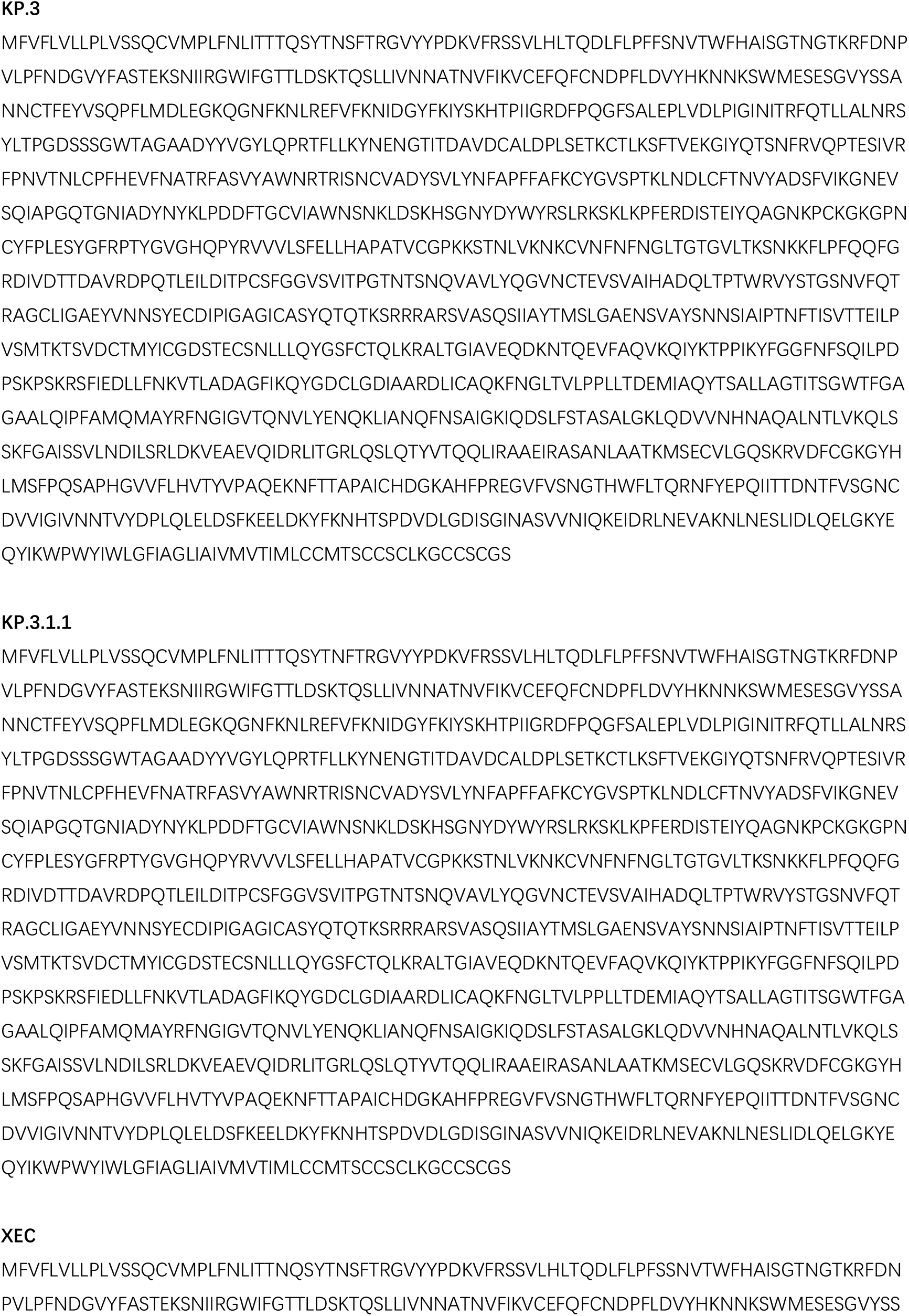
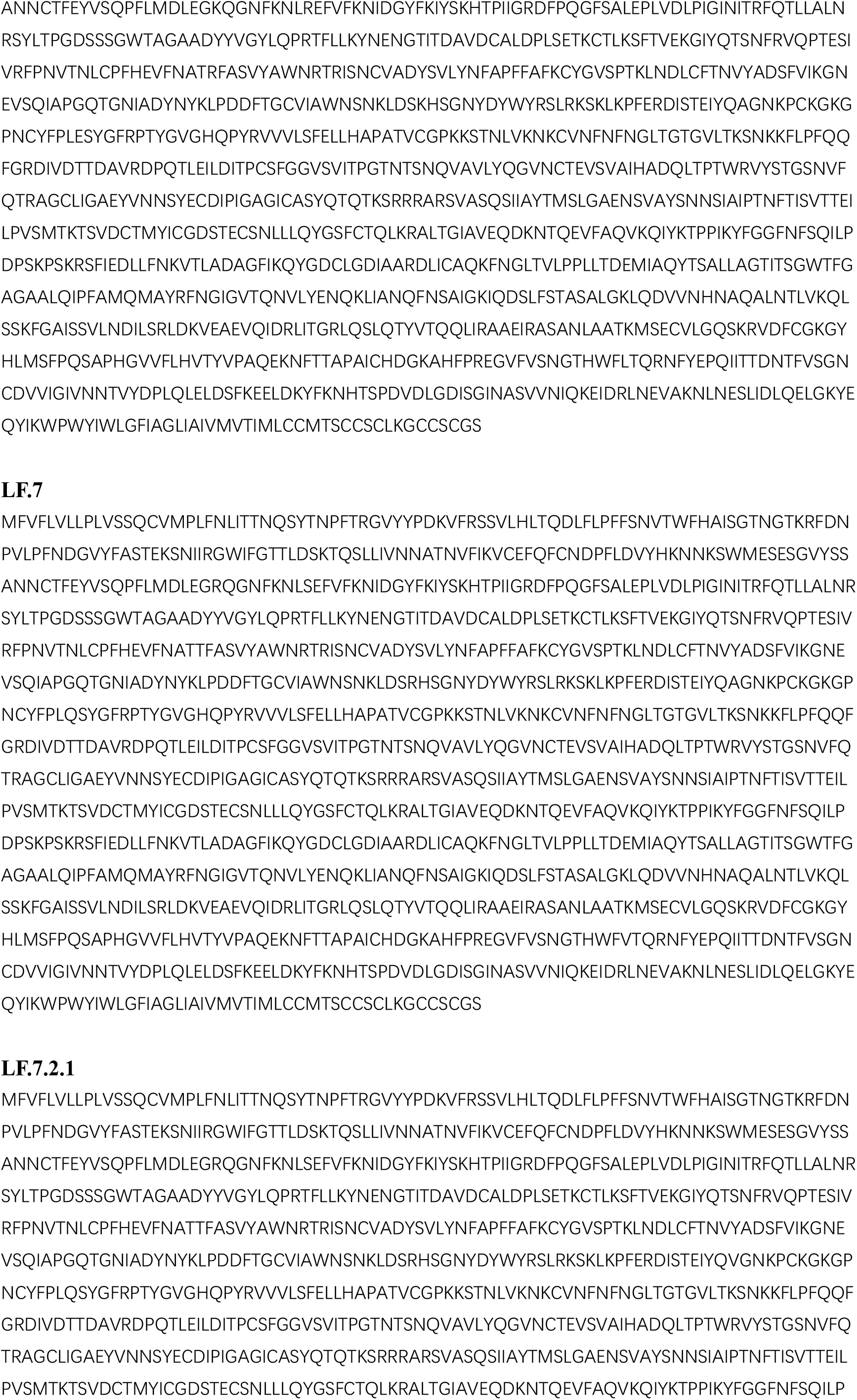
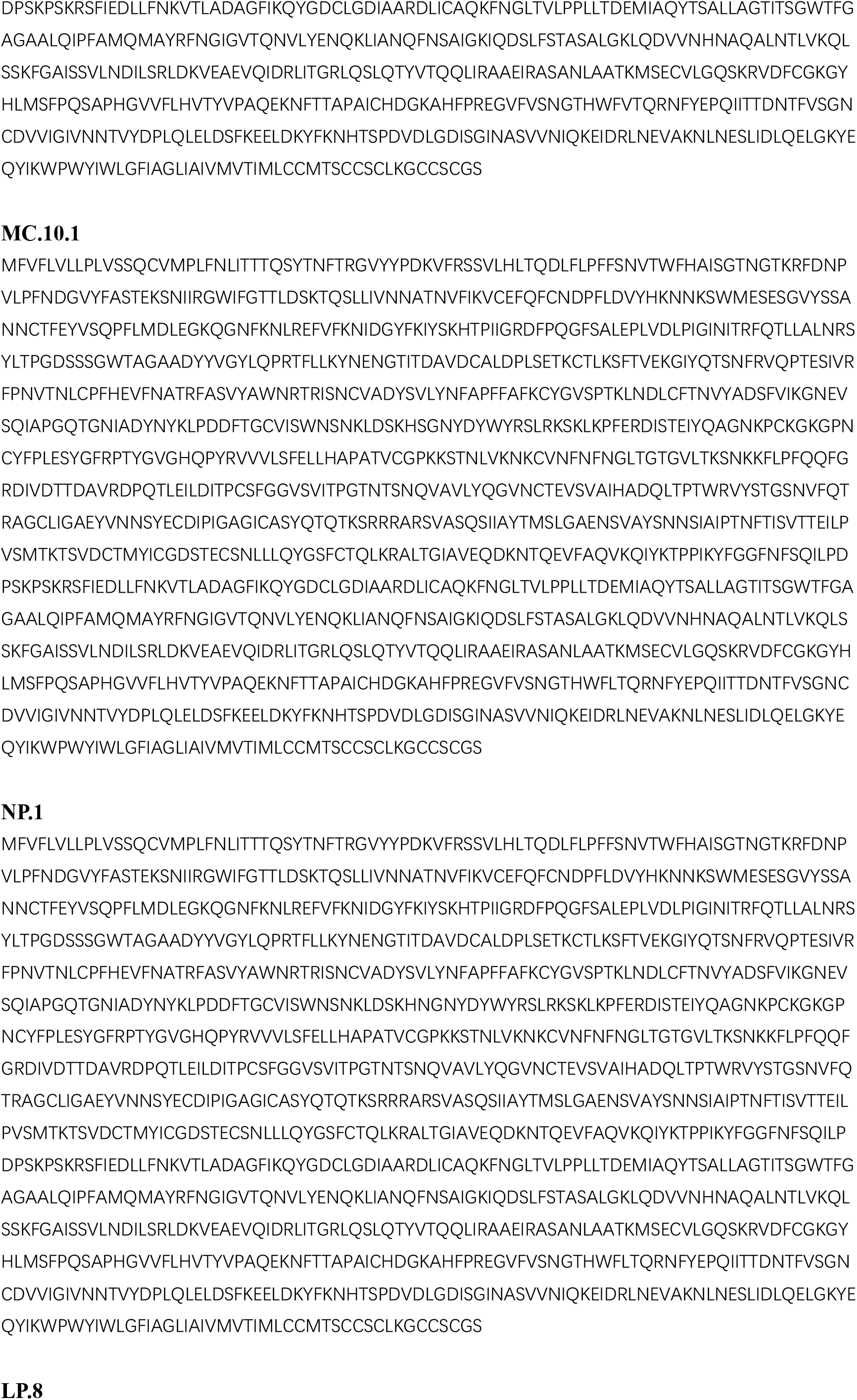
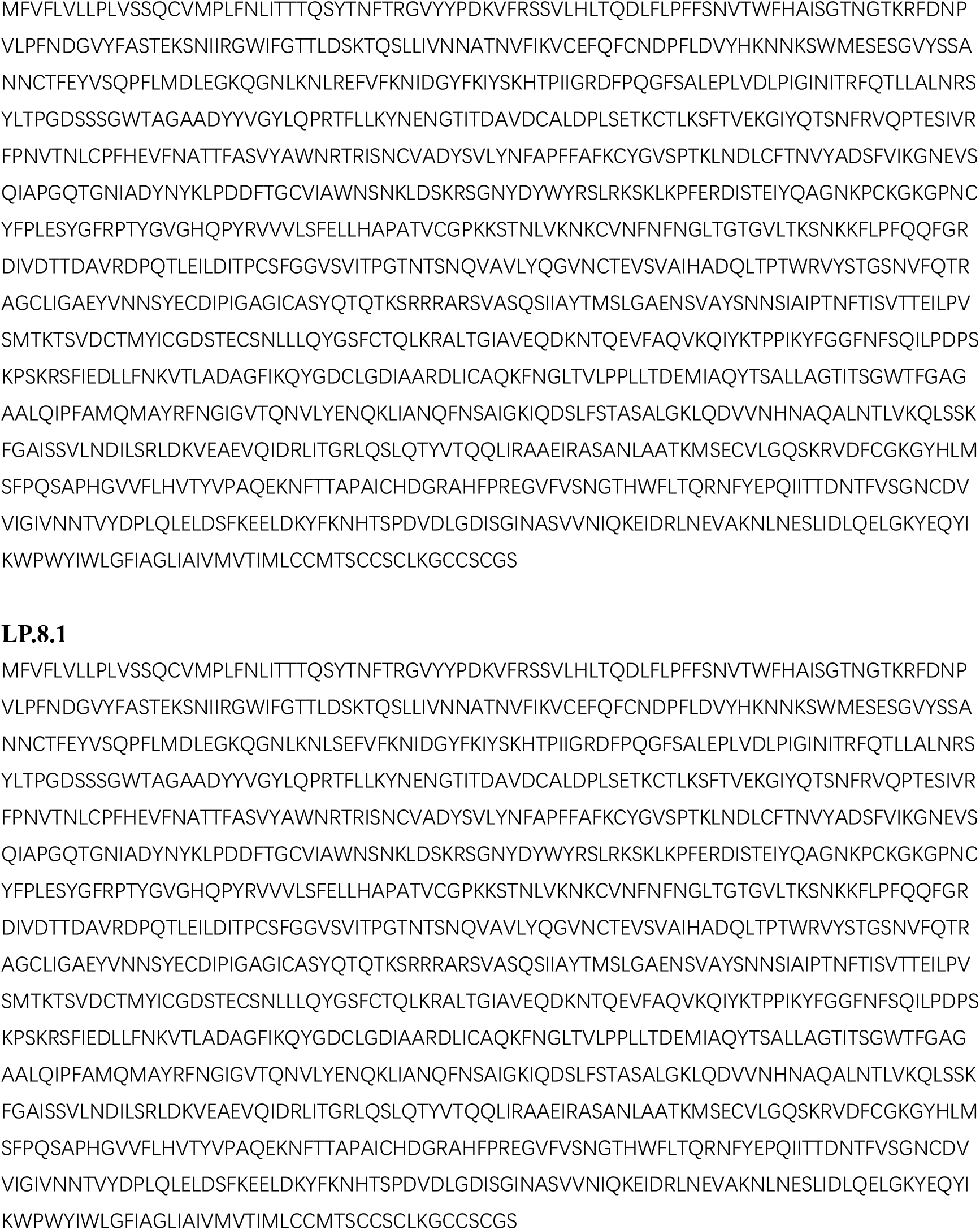
Spike protein sequences of SARS-CoV-2 variants.

**Table S2.**
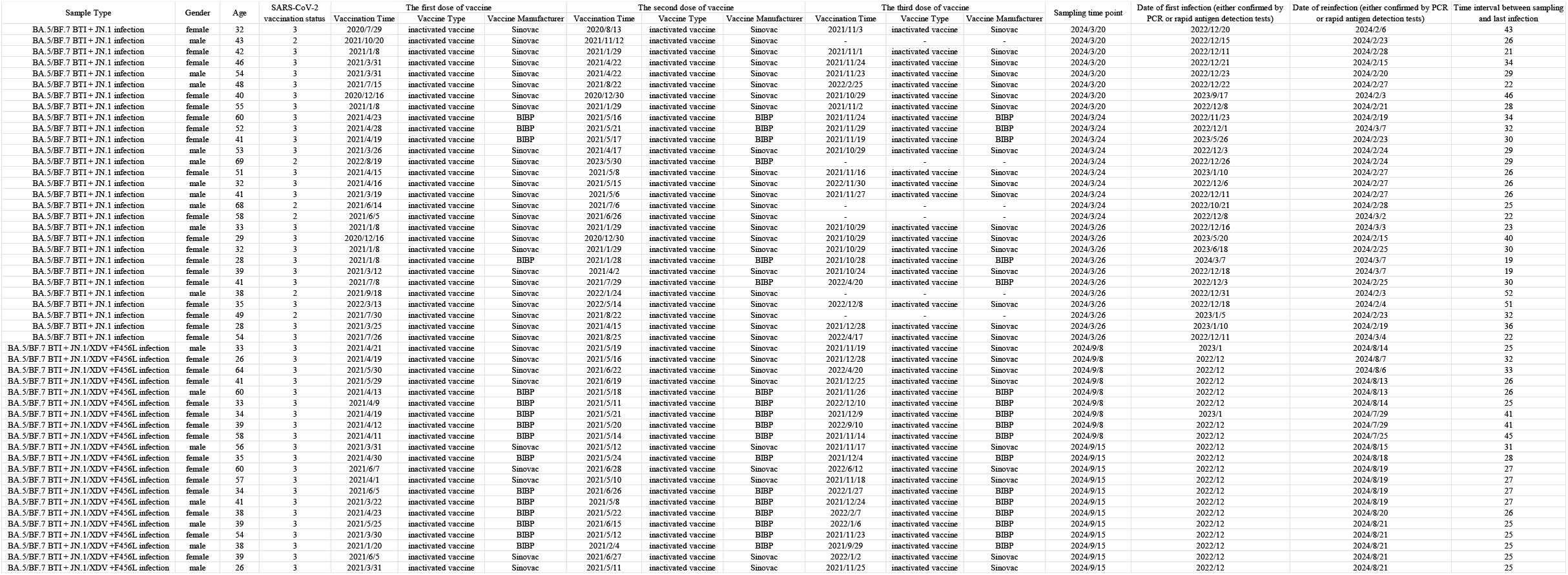
Information of SARS-CoV-2 convalescent patients involved in the study.

